# Disparate co-evolution and prevalence of sulfadoxine and pyrimethamine resistance alleles and haplotypes at *dhfr* and *dhps* genes across Africa

**DOI:** 10.1101/2024.12.16.628681

**Authors:** Nina F. D. White, Georgia Whitton, Varanya Wasakul, Lucas Amenga-Etego, Antoine Dara, Voahangy Andrianaranjaka, Milijaona Randrianarivelojosia, Olivo Miotto, Umberto D’Alessandro, Abdoulaye Djimdé, Cristina V. Ariani, Richard D. Pearson, Alfred Amambua-Ngwa

## Abstract

Sulfadoxine-pyrimethamine (SP), despite the emergence and spread of mutations in *dhfr* and *dhps* genes associated with lower treatment efficacy, is still recommended alone or in combination by the WHO for preventive treatment in pregnant women, children and infants. Therefore, it is important to understand the evolution of *P. falciparum dhfr* and *dhps* genes. Here, we used a subset of the MalariaGEN Pf7 dataset to describe haplotype frequencies across 22 African countries, including changes over time in The Gambia, Mali, Ghana and Kenya. We show that the triple mutant of *dhfr,* N51I/C59R/S108N, has remained the dominant haplotype across the continent with limited evidence of additional mutations. There is greater variation for *dhps*, with a total of 51 different haplotypes present. The *dhps* resistance mutation A437G has risen close to fixation across most of Africa, although at a lower frequency in the northwest malaria-endemic part of the continent (Gambia, Senegal and Mali). The A437G mutation is usually found together with K540E in East Africa, but K540E is still very rare in West Africa. Although samples from Madagascar have low genetic differentiation from samples from mainland East Africa at the whole genome level, we show that *dhps* K540E is highly differentiated between the two populations, being at very low frequency in Madagascar (4%). We used whole genome data to show that only 12 SNPs are more highly differentiated than K540E between Madagascar and East Africa, with *aat1* and a possible novel drug resistance locus approximately 20kb 3’ of *mdr1* having even higher F_ST_. We highlight the value of longitudinal sampling and whole genome sequence data for understanding the heterogeneity and ongoing changes in anti-malarial drug resistance genetic markers.

## Introduction

As malaria continues to cause significant morbidity and mortality, especially in Africa, monitoring of current interventions and implementation of new ones are needed. Malaria particularly affects children and pregnant women, who contributed to most of the over 240 million infections and 600,000 deaths reported in 2022 (WHO, 2023b). To prevent morbidity and mortality in these vulnerable groups, malaria-endemic countries in Africa are employing Sulfadoxine-Pyrimethamine (SP) in various methods. SP is used for intermittent preventive treatment during pregnancy (IPTp), for perennial malaria chemoprevention (PMC) in infants or, in combination with amodiaquine (AQ) for seasonal malaria chemoprevention (SMC) in children 3-59 months old. SP is deployed across 15 countries in the Sahel region of West Africa, and in Mozambique and Uganda in East Africa (WHO, 2023a). SP has been previously used as the first-line treatment for malaria. In addition, AQ is a partner drug in one of the currently available artemisinin-based combination therapies (ACTs) to treat patients with uncomplicated malaria. Although SP-AQ remains effective for chemoprevention, resistance to SP and AQ separately has long been established, with SP discontinued as a first-line therapy since the early 2000s in most endemic countries (World Health Organization, 2006). Hence the continuous use of SP in IPTp, PMC and SMC may further select resistant *P. falciparum* strains that could render the drugs even less effective in the future.

Resistance to SP has been attributed to mutations in *P. falciparum* dihydrofolate reductase (*dhfr*) and dihydropteroate synthetase (*dhps*) genes for pyrimethamine and sulfadoxine components, respectively (Plowe *et al*., 1997). In African malaria parasite populations, resistance-associated haplotypes in these genes were imported from Asia and spread by transmission (Roper *et al*., 2004; Mita *et al*., 2011). For *dhfr*, the following mutations; N51I, C59R, S108N, and I164L form the resistant haplotype, with IRN at 51, 59, and 108 reported at high frequencies across the African continent (Sridaran *et al*., 2010). Sulfadoxine resistance-associated haplotypes of *dhps* at positions S436A/F, A437G, K540E, A581G, and A613S/T are more complex, with heterogeneous combinations and distribution across the African continent (Sridaran *et al*., 2010; Amimo *et al*., 2020). SP resistance results from different combinations of mutations across both the *dhfr* and *dhps* genes. For example, the quintuple mutant, a combination of *dhfr*-IRN and *dhps*-GE, confers a larger risk of treatment failure than the individual *dhfr* triple or *dhps* double mutants (Happi *et al*., 2005). Beyond these, *dhfr*-I164L and *dhps*-A581G mutations have been associated with a higher risk of treatment failure in East Africa (Mbogo *et al*., 2014; Juliano *et al*., 2024). Alongside *dhps*-A613S/T, these two mutations in combination with the quintuple mutant are said to confer ‘super-resistance’ to SP (Naidoo and Roper, 2013). The prevalence of both rare and common haplotype combinations, and their effect on chemoprevention is relevant for strategizing drug intervention policies toward control and elimination.

Malaria Molecular Surveillance (MMS) of malaria parasites for markers of resistance and diversity can complement current epidemiological approaches against malaria and inform the optimal deployment of interventions. Several studies have employed microsatellite and SNP typing approaches to map the prevalence, origins, and dispersal of SP-resistance markers across Africa and the globe. In the last decade, with advances in genome sequencing of *P. falciparum*, the Pathogen (Plasmodium) Diversity Network Africa (PDNA) consortium in collaboration with MalariaGEN embarked on genomic sequencing of malaria parasites across 20 African countries. This report presents the prevalence of genetic markers of SP resistance across both PDNA sites and other African countries. It shows the different rates and directions of selection of these alleles over 30 years in four African countries, especially for the *dhps* markers of sulfadoxine resistance

## Results

Whole-genome sequences from the MalariaGEN Pf7 dataset were used in this study (MalariaGEN *et al*., 2023). Pf7 contains 8,492 QC-passed *Plasmodium falciparum* samples from 22 African countries which were selected for analysis. Amino acid haplotypes for *dhfr* and *dhps* were determined for each sample by applying all mutations discovered in the sample to the 3D7 reference sequence, then translating the nucleotide sequence of concatenated exons. Frequencies of individual mutations over time were generated using the allele frequency method, which considers only homozygous samples in the calculation (see Box).

### Variation in *dhfr*

The most frequent haplotype of *dhfr* was the triple mutant (N51I/C59R/S108N), found in 80% of samples (Figure 1, Supplementary Table S1). The second most frequent haplotype, found in 10% of samples, was the 3D7 reference genome sequence - which is thought to be the wild type for *dhfr*. The third and fourth most common haplotypes were C59R/S108N and N51I/S108N, respectively. These two haplotypes had distinct geographical spreads, C59R/S108N was more common in West and East Africa, while N51I/S108N was more common in Central and North-East Africa (Figure 1). The other potential combination of the N51I, C59R, and S108N mutations, N51I/C59R, was not observed. All three mutations N51I, C59R, and S108N were seen in isolation (Supplementary Table S1). However, the N51I-only and C59R-only haplotypes were rare, each recorded in only two and one sample(s), respectively. The S108N-only haplotype was more common, being found in 25 samples (0.4%). The I164L mutation was very rare in this dataset and seen only twice: once together with the N51I/C59R/S108N triple mutant and once with N51I and S108N (Figure 2b, Supplementary Table S1). In addition to the five most common haplotypes shown in Figure 1, there were 14 additional *dhfr* haplotypes, each seen in either one or two samples, and having a combined frequency of 0.25% (Supplementary Table S1).

**Figure 1.**
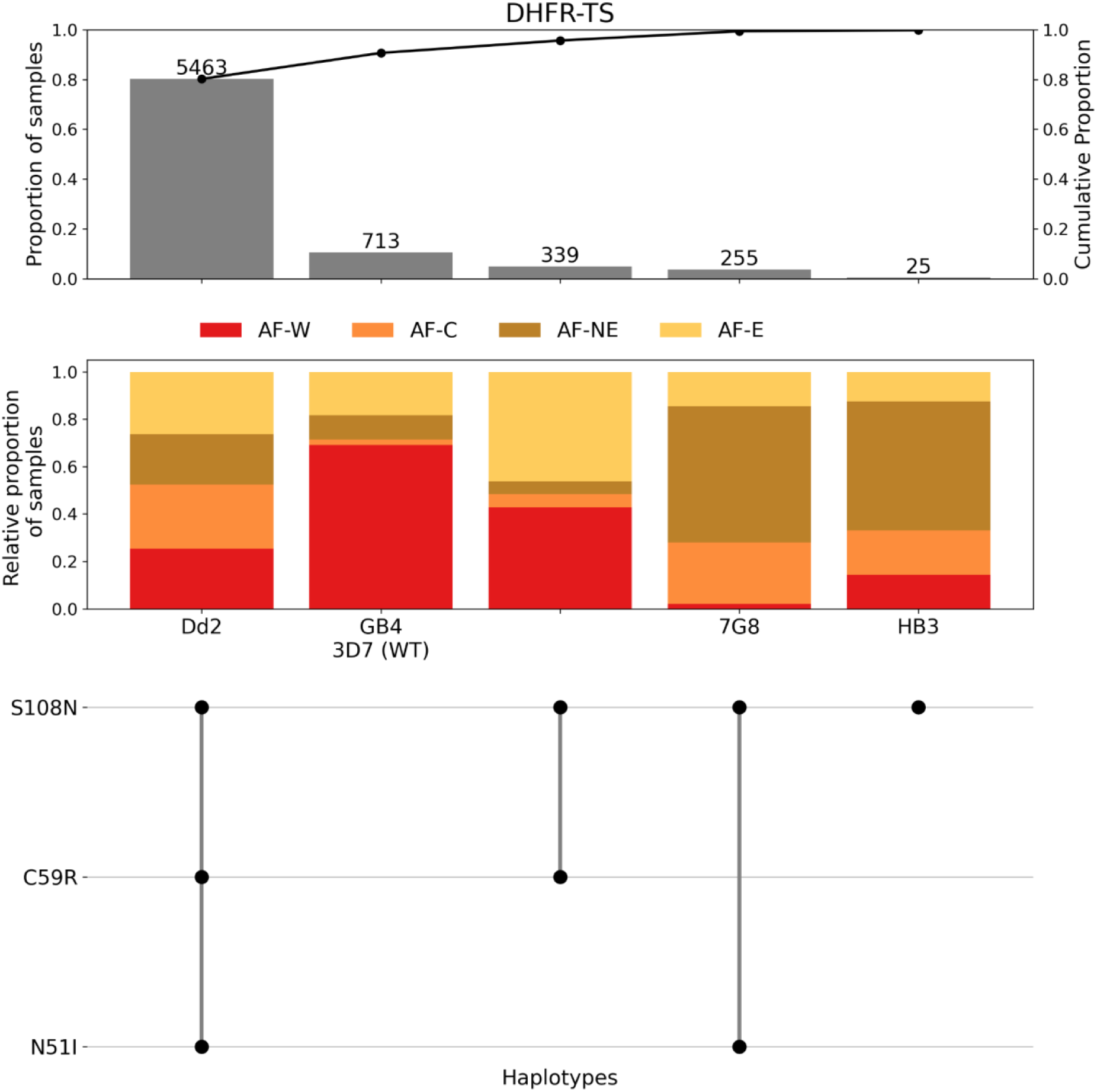
Summary of the five most frequent *dhfr* haplotypes across 21 African countries. The frequency of each haplotype, in terms of sample count (number above each bar), frequency across the dataset (1^st^ y-axis), and cumulative proportion of samples (2^nd^ y-axis) is shown in the first panel. The proportion of each haplotype assigned to each of the major African subpopulations is shown in the second panel (AF-W = West Africa, AF-C = Central Africa, AF-NE = North-East Africa, AF-E = East Africa). The final panel illustrates the individual amino acid mutations contributing to each unique haplotype. The 3D7 wild type (WT) haplotype contains no mutations, so remains blank in the final panel.

**Figure 2.**
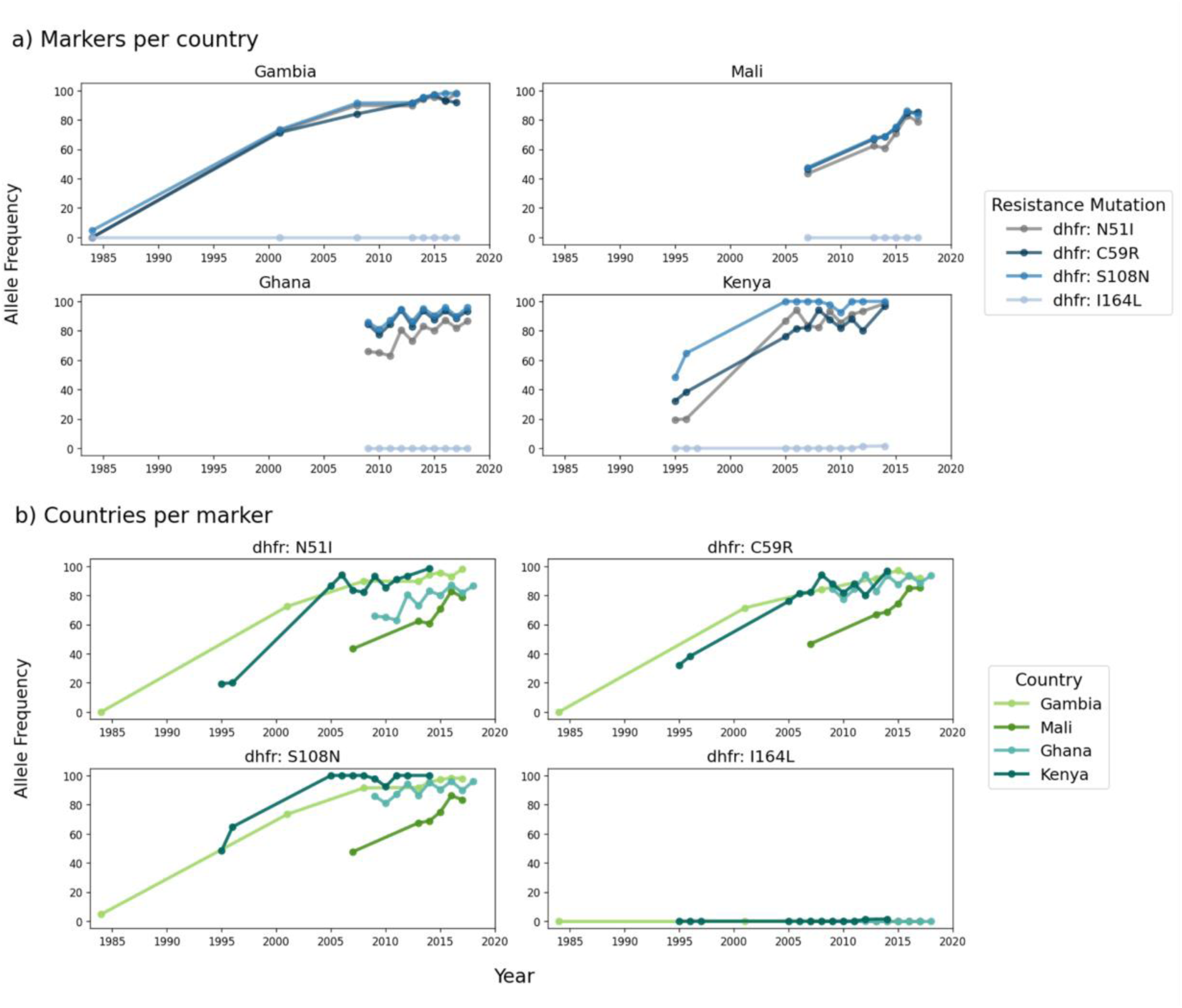
Temporal analysis of *dhfr* allele frequencies. a) All *dhfr* mutations within Gambia, Mali, Ghana, and Kenya, b) comparison of allele frequencies for individual *dhfr* mutations across countries. Lines represent the allele frequency of that mutation originating from homozygous genotypes only. Allele frequencies were calculated only for years with at least 25 homozygous samples (see Box for further details).

For many of the countries, there were only data from one or two different years, making it impossible to determine trends. Time series analysis was possible for The Gambia, Mali, Ghana and Kenya as there were multiple time points (Figure 2). All three mutations, i.e., N51I, C59R, and S108N, increased in frequency over time (Figure 2a). In Kenya, S108N rose in frequency earlier than N51I and C59R, as expected given the established step-wise accumulation of these three *dhfr* mutations that these latter mutations have occurred on an S108N background in this East African setting. However, in West African countries, these three mutations increased in frequency at approximately the same rate. The prevalence of S108N increased earlier in Kenya, followed by The Gambia and Ghana, and then Mali (Figure 2b). N51I prevalence also increased earlier in Kenya and The Gambia, followed by Ghana and then Mali. The same trend was observed for C59R.

### Variation in *dhps*

Compared with *dhfr*, there was a much greater diversity of haplotypes in *dhps*. The eight most frequent *dhps* haplotypes were all seen at a frequency greater than 1%, while there were a further 43 *dhps* haplotypes with a combined frequency of 3.5% (Figure 3, Supplementary Table S2). Over half of these (24) were singleton haplotypes seen in only one sample (Supplementary Table S2). The most frequent *dhps* haplotypes showed distinct geographic patterns (Figure 3). Six of the eight most common haplotypes contained the mutation A437G (Figure 3). Four haplotypes with A473G and without K540E were found predominantly in West and Central Africa. The single mutation haplotype A437G was most common of all *dhps* haplotypes, found in 36% of all samples. Two haplotypes containing A437G alongside K540E were restricted to the other side of the continent, in East and Northeastern Africa (Figure 3). A437G/K540E was the dominant haplotype in East Africa (74%, 907/1,234) and found in (18% of all African samples) (Figure 3). Of the eight most common *dhps* haplotypes, two did not contain A437G (Figure 3). These were both more geographically widespread relative to the other six common haplotypes. The single mutation haplotype S436A, was found in 11% of samples and occurred in all four African populations from this study (though predominantly was found in West Africa). The wild type *dhps* sequence, with no mutations, was the fifth most common haplotype, being seen in 7.8% of samples (Figure 3). Of note is that the reference 3D7 strain does not contain the wild type haplotype for *dhps,* though the lab strains HB3 and IT have this haplotype. The wild type was most evenly distributed geographically of the eight most common *dhps* haplotypes.

**Figure 3.**
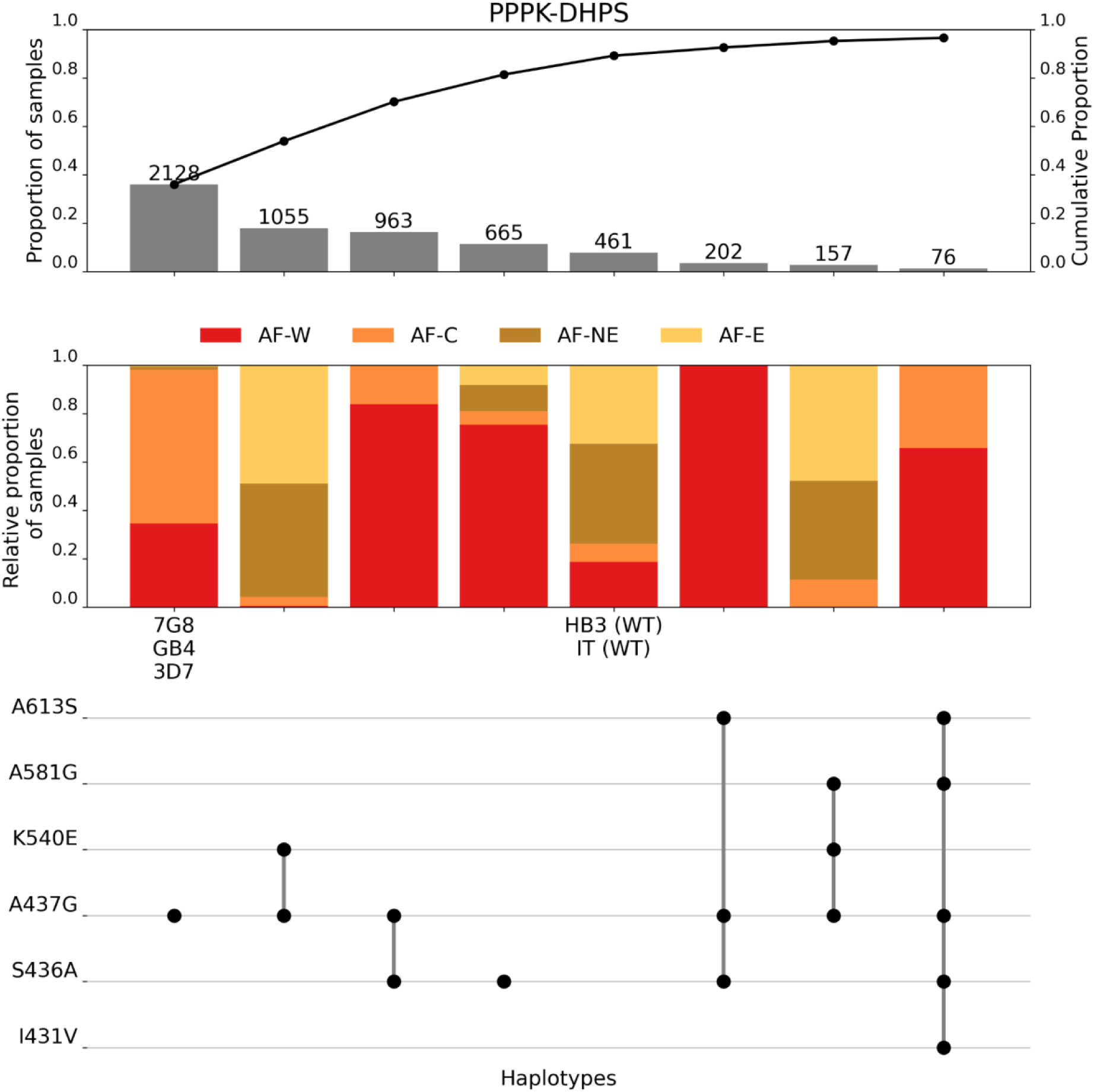
Summary of the eight most frequent *dhps* haplotypes across 21 African countries. The frequency of each haplotype, in terms of sample count (number above the bar), frequency across the dataset (1^st^ y-axis), and cumulative proportion of samples (2^nd^ y-axis) is shown in the first panel. The proportion of each haplotype assigned to each of the major African subpopulations is shown in the second panel (AF-W = west Africa, AF-C = central Africa, AF-NE = northeast Africa, AF-E = east Africa). The final panel illustrates the individual amino acid mutations contributing to each unique haplotype. The HB3 and IT strains represent the wild type (WT) haplotype, so no mutations are shown here.

We have data from samples collected in the 20th century from both the Gambia and Kenya. In both cases, the frequency of the A437G mutation has increased since pre-2000 (Figure 4). A437G reached near fixation in Kenya (2014) and a high of 80% in The Gambia. Between 2007 and 2017, A437G prevalence in Mali increased, while in Ghana it remained at a stable, high frequency between 2009 and 2018. Spatially, A473G has been more common in the south and east of West Africa than in the north and west (Figure 5a). The frequency of the S436A mutation has increased over time in The Gambia but has fallen slightly in both Mali and Ghana over time (Figure 4b). The frequency of S436A has remained low in Kenya over time.

**Figure 4:**
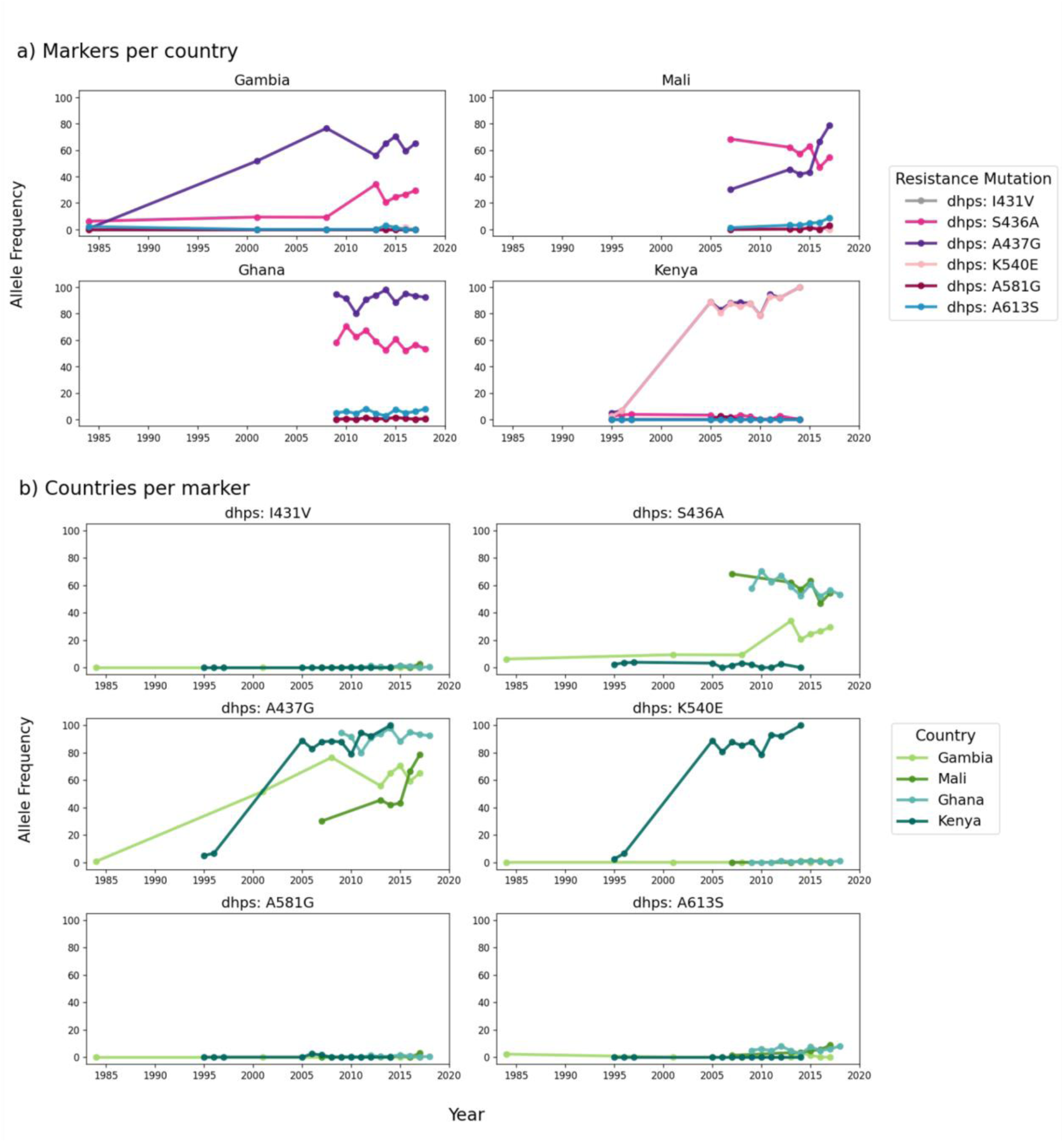
Temporal analysis of *dhps* allele frequencies. a) all *dhps* mutations within Gambia, Mali, Ghana, and Kenya are shown in the first four plots, b) comparison of allele frequencies for individual *dhps* mutations across countries. Lines represent the allele frequency of that mutation originating from homozygous genotypes only. Allele frequencies were calculated only for years with at least 25 homozygous samples (see Box for further details).

**Figure 5:**
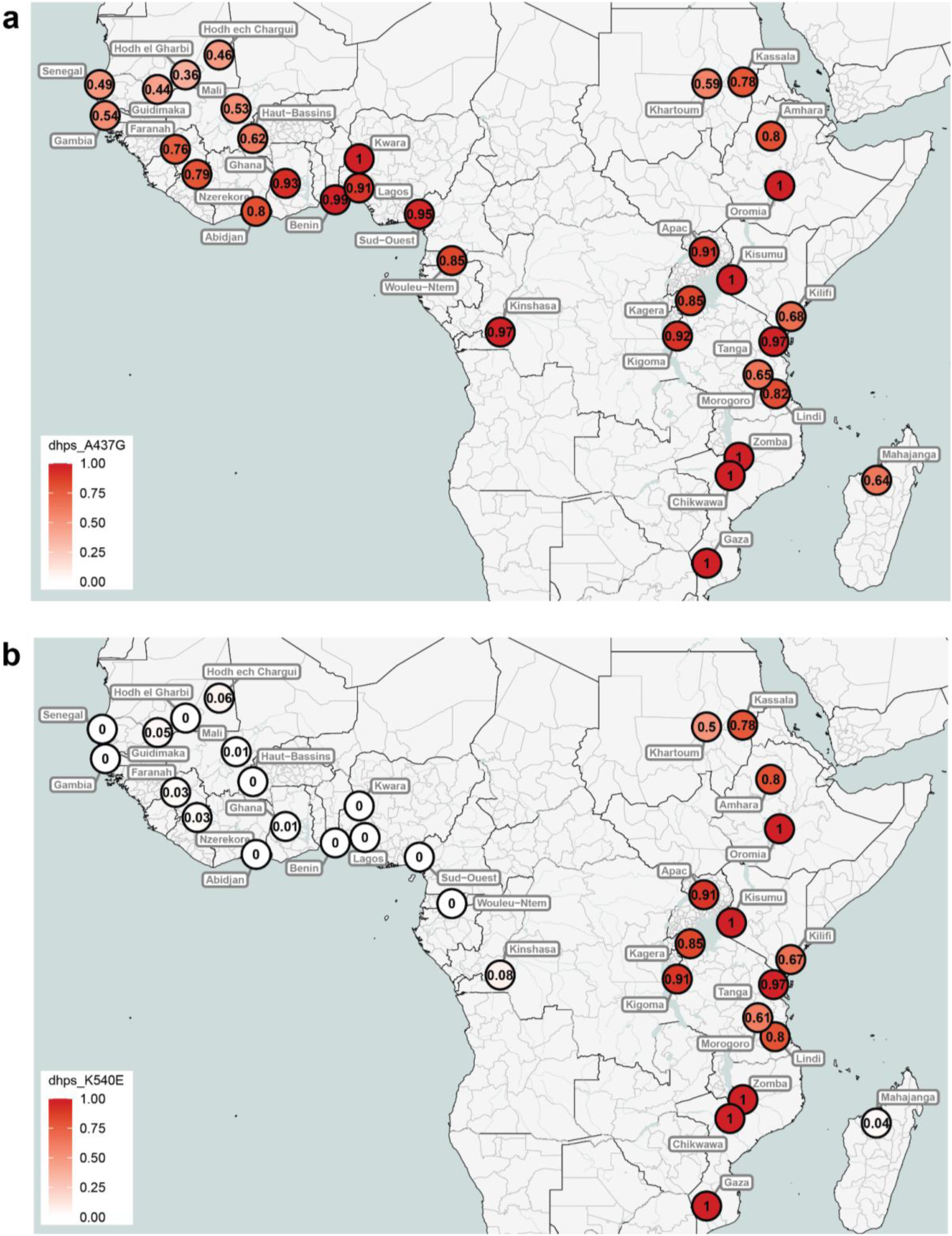
Distribution of mutation frequencies for a) dhps A437G and b) dhps K540E. Data were aggregated at the administrative division 1 level. To improve visualisation and avoid overlapping markers for nearby locations, data from Senegal, Gambia, Ghana, Mali and Benin were combined at the country level for panel b. Colours represent mutation frequencies ranging from 0 (no mutation detected) to 1 (100% of the parasites in the location carrying the mutation).

Of the four countries for which we have time series data, the K540E mutation is seen almost exclusively in Kenya, where it reached close to fixation in recent years (Figure 4b). It is interesting to note that in Kenya, the frequency of K540E has tracked almost exactly the frequency of A437G, suggesting co-selection in this geographical region. Although K540E is widespread across East Africa, it is very rare in both West Africa and Madagascar (Figure 5b). Interestingly, despite such a high differentiation at K540E between East Africa and Madagascar, these populations are similar at a whole genome level (Supplementary Figure S1). It is important to note that data from only 24 isolates from Madagascar were available for this analysis.

To further investigate the extent of differentiation at K540E between East Africa and Madagascar, pairwise F_ST_ values for QC-passed, genome-wide SNPs were generated between samples from these two subpopulations. Values across the genome ranged between 0 and 0.91, with a median of 0.0003 across all genome-wide variants (Table 1). The F_ST_ value for K540E was 0.748, which was greater than 99.99% of genomic positions analysed, with just 12 positions exhibiting a greater difference than 0.748 (Table 1, Supplementary Figures S2 and S3). Note that F_ST_ could be calculated for 763,535 positions. When investigating the 20 most differentiated SNPs between mainland East Africa and Madagascar, we observed SNPs encoding non-synonymous amino acid mutations within *aat1* and approximately 20kb 3’ of *mdr1* which had even higher F_ST_ than the SNP encoding K540E (Table 1). These may suggest novel drug-resistance mutations.

**Table 1:** The 20 most differentiated single nucleotide polymorphisms between AF-E (n = 1,515) and Madagascar (n = 24), based on alternate allele frequency. REF = 3D7 reference strain nucleotide, ALT = alternate nucleotide.

| Genomic Position | REF | ALTs | East Africa<br>ALT Freq. | Madagascar<br>ALT Freq. | F<sub>ST</sub> | Gene ID | Gene Name | Amino Acid<br>Change | Mutation Type |
| --- | --- | --- | --- | --- | --- | --- | --- | --- | --- |
| Pf3D7_05_v3:1141578 | A | C | 0.00 | 0.92 | 0.91 | Pf3D7_0527400 |  |  | 3' UTR variant |
| Pf3D7_05_v3:1145481 | C | A | 0.00 | 0.92 | 0.91 | Pf3D7_0527500 | HIP |  | 3' UTR variant |
| Pf3D7_05_v3:1139023 | A | G | 0.01 | 0.92 | 0.90 | Intergenic region: Pf3D7_0527300 -<br>Pf3D7_0527400 |  |  | Intergenic region |
| Pf3D7_05_v3:1140984 | T | G | 0.00 | 0.90 | 0.89 | Pf3D7_0527400 |  | Leu120Val | Non-synonymous |
| Pf3D7_05_v3:1140987 | G | A | 0.00 | 0.90 | 0.89 | Pf3D7_0527400 |  | Asp121Asn | Non-synonymous |
| Pf3D7_05_v3:1140988 | A | G | 0.00 | 0.90 | 0.89 | Pf3D7_0527400 |  | Asp121Gly | Non-synonymous |
| Pf3D7_06_v3:1215233 | G | A | 0.87 | 0.00 | 0.87 | Pf3D7_0629500 | AAT1 | Ser258Leu | Non-synonymous |
| Pf3D7_06_v3:1214259 | C | G | 0.84 | 0.00 | 0.84 | Pf3D7_0629500 | AAT1 |  | Intron variant |
| Pf3D7_06_v3:1214350 | G | T | 0.86 | 0.04 | 0.80 | Pf3D7_0629500 | AAT1 | Ile552Ile | Synonymous |
| Pf3D7_05_v3:1151093 | C | A | 0.00 | 0.81 | 0.81 | Intergenic region: Pf3D7_0527700 -<br>Pf3D7_0527800 |  |  | Intergenic region |
| Pf3D7_08_v3:546841 | T | A | 0.78 | 0.00 | 0.78 | Pf3D7_0810700 |  | Lys68Asn | Non-synonymous |
| Pf3D7_05_v3:1133912 | T | A | 0.00 | 0.77 | 0.76 | Pf3D7_0527200 | USP14 | Thr281Thr | Synonymous |
| Pf3D7_08_v3:549993 | A | G | 0.81 | 0.04 | 0.75 | Pf3D7_0810800 | PPPK-DHPS | Lys540Glu | Non-synonymous |
| Pf3D7_05_v3:1119264 | A | G | 0.82 | 0.08 | 0.70 | Pf3D7_0526700 |  | Tyr130His | Non-synonymous |
| Pf3D7_14_v3:2183418 | A | T | 0.05 | 0.79 | 0.72 | PF3D7_1453200 |  | Asn756Tyr | Non-synonymous |
| Pf3D7_05_v3:1114069 | C | T,G | 0.07 | 0.79 | 0.69 | PF3D7_0526600 |  | Ile4289Met (>G),<br>Ile4289Ile (>T) | Non-synonymous (>G),<br>Synonymous (>T) |
| Pf3D7_14_v3:2183653 | C | A | 0.01 | 0.73 | 0.70 | PF3D7_1453200 |  | Ser834Tyr | Non-synonymous |
| Pf3D7_14_v3:2183719 | G | A | 0.05 | 0.77 | 0.70 | PF3D7_1453200 |  | Arg856Gln | Non-synonymous |
| Pf3D7_06_v3:1206498 | G | A | 0.71 | 0.00 | 0.69 | PF3D7_0629300 | PL | Glu437Lys | Non-synonymous |
| Pf3D7_06_v3:1201098 | T | C | 0.69 | 0.00 | 0.69 | PF3D7_0629200 | DnaJ |  | 5' UTR variant |

### Reliability of using single markers as proxies for full gene haplotypes

It has been suggested that the order in which mutations accumulated in *dhfr* was S108N, followed by N51I, followed by C59R (Lozovsky *et al*., 2009; Brown *et al*., 2010). If this were indeed the case, then it would be possible to use the marker C59R as a proxy for the N51I/C59R/S108N triple mutant (Supplementary Table S3). This study demonstrates that there are examples when, for example, C59R is seen in haplotypes that include S108N but not N51I. Therefore, C59R is certainly not a perfect proxy for the triple mutant. The reliability of the C59R marker as a proxy for the triple mutation can be assessed by determining what proportion of samples containing this mutation do indeed have the triple mutation. In all countries, this reliability score is over 80% (Figure 6). However, reliability differs across countries, for example being only 85% in Kenya. If positions 51, 59 and 108 are genotyped, the vast majority of variation will be captured. Given that mutations at positions 51 and 59 are almost always found on the background of S108N, genotyping one, or both, of them should be sufficient to determine most of the variation within the gene.

**Figure 6.**
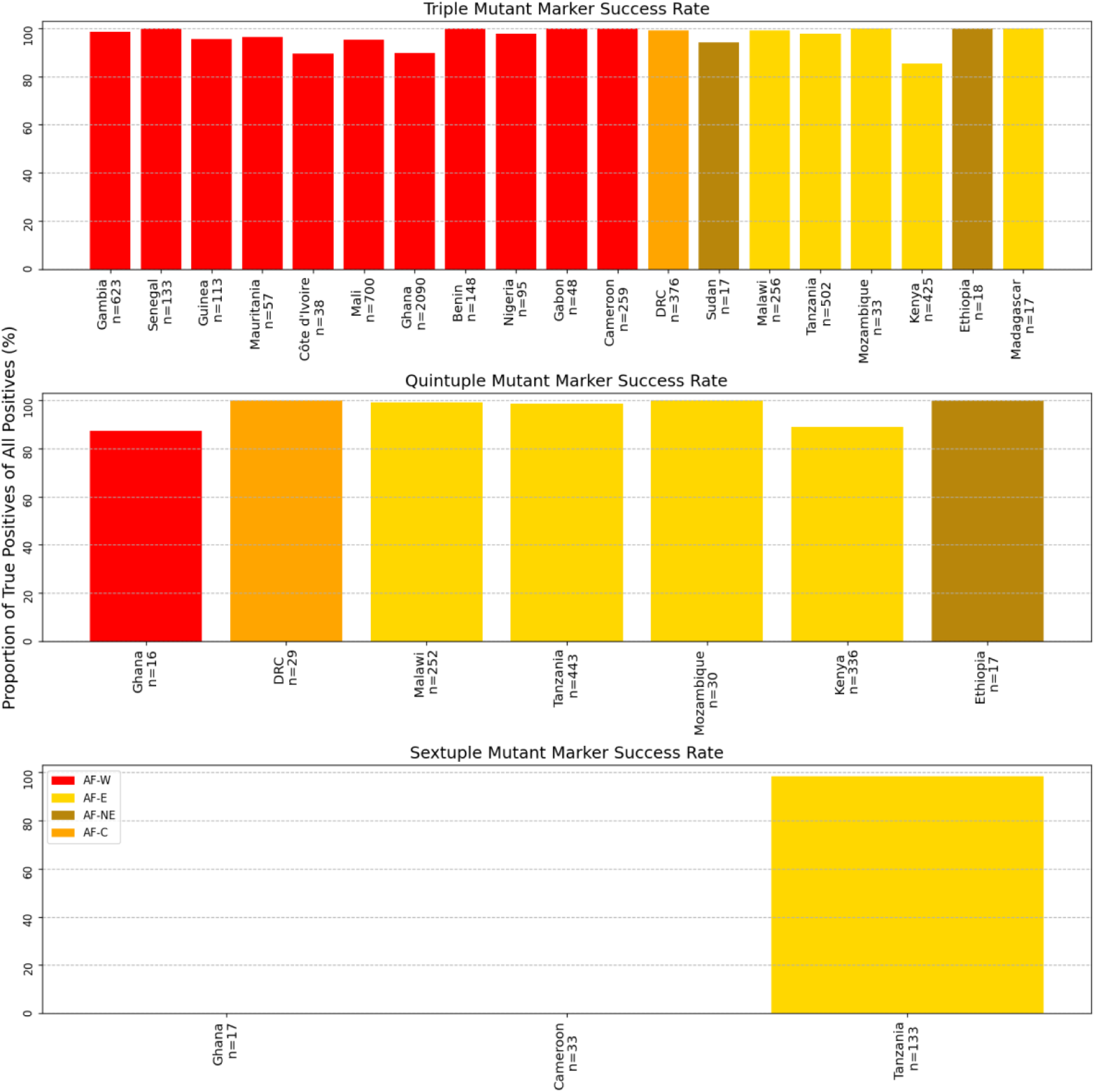
Reliability of single mutation markers of haplotypes. Success rate of suggested markers to genotype for the triple, quintuple and sextuple multi-mutant haplotypes in all African countries where n > 15. Triple = *dhfr* C59R for *dhfr* C59R/N51I/S108N. Quintuple = *dhfr* C59R and *dhps* K540E for *dhfr* C59R/N51I/S108N, *dhps* A437G/K540E. Sextuple = dhps581[G] for *dhfr* N51I/C59R/S108N, *dhps* A437G/ K540E/A581G. Bars are coloured according to major subpopulation (AF-W = west Africa, AF-C = central Africa, AF-NE = northeast Africa, AF-E = east Africa).

Likewise, the combination of C59R and K540E markers are generally reliable markers of the quintuple multi-gene haplotype *dhfr* N51I/C59R/S108N, *dhps* A437G/K540E. However, the reliability of this combination of markers is lower in Kenya (89%) than in many other countries (Figure 6). Finally and based on our results, while in Tanzania A581G seems to be a reliable marker of the multi-gene, sextuple haplotype *dhfr* N51I/C59R/S108N and *dhps* A437G/K540E/A581G, this was not the case across Africa. A581G in West African countries such as Ghana and Cameroon was never a proxy for this sextuple haplotype (Figure 6). Therefore, a marker may be a reliable proxy of a haplotype in one region but not necessarily in other regions.

## Discussion

SP resistance emerged approximately three decades ago and led to the discontinuation of SP as the sole first-line treatment for malaria in the early 2000s (World Health Organization, 2006). However, SP is used for chemoprevention during pregnancy, infancy and childhood. Thus, monitoring genetic markers in *dhfr* and *dhps* associated with SP resistance is informative for interventions relying on SP. This study describes the evolution of mutation associated to SP resistance across Africa, the continent with the highest malaria burden, where SP-based interventions are widely implemented.

Prevalence of the triple mutant *dhfr* haplotype was high across Africa, present in over 80% of all samples analysed. The prevalence of the mutations *dhfr* N51I, C59R, and S108N in The Gambia, Ghana, Mali, and Kenya increased over 30 years to near-fixation, possibly due to the selection exerted by the widespread use of SP (Sridaran *et al*., 2010). Despite the withdrawal of SP as first-line treatment and thus a decreased evolutionary pressure over the last 20-25 years, the parasite wild types did not increase. Rather, the high prevalence of the *dhfr* mutations suggest an ongoing and continuous selection for SP resistance, with the risk of impacting on the SP-based interventions.

Although the selective pressure exerted by the SP used for IPTp, PMC or SMC may explain in some countries the high prevalence of the triple mutant, this could also be due to the acquisition of other mutations compensating the loss of fitness associated to the mutations. This is supported by the low but significant prevalence of haplotypes combining two of the three *dhfr* triple mutant loci. These haplotypes were C59R/S108N and N51I/S108N and were the third and fourth most common *dhfr* haplotypes, respectively. The accumulation of the triple mutant alleles seen in this study is concordant with the widely accepted step-wise order of 108, then N51I and C59R (Plowe *et al*. 1997). In Kenya, the earlier fixation of S108N supports the theory of later introduction of N51I and C59R. In the Gambia, however, all three triple mutant alleles increased at the same rate, not in contrast to the stepwise accumulation hypothesis, but also not clearly demonstrating the event.

There was no joint N51I/C59R *dhfr* haplotype, suggesting that both N51I and C59R have arisen on an S108N background, which is the most frequent single mutant of *dhfr*. The N51I/C59R and C59R-only haplotypes have been described in Melanesia, and are thought to have originated independently from the dominant N51I/C59R/S108N triple mutant haplotype (Wang *et al*., 1997). The *dhfr* C59R-only haplotype was first reported in Vietnam, though at low frequencies (Mita *et al*., 2007). In Central and North-East Africa, C59R/S108N was rare, while N51I/S108N was found at over 25% in these populations. Therefore, N51I and C59R may have been introduced, emerged, or been selected differently in different populations on the S108N background. Selection of independently emerging double mutants has been shown in previous studies in Ghana and Kenya, with non-Southeast Asian indigenous haplotypes of double mutants on the S108N background (Mita, Tanabe and Kita, 2009). Both double haplotypes (N51I/S108N and C59R/S108N) in West and East Africa may have spread by recent human migration or may be due to the breakdown of triple haplotype by recombination or reverse mutation. However, this is extremely difficult to establish. S108N was probably the earliest mutation in Africa, followed by multiple introductions of either or both N51I and C59R from Southeast Asia. Various studies of the fitness and epidemiology of combined mutations have suggested a stepwise evolution of mutants; NCNI → ICNI or NRNI → IRNI → IRNL (Lozovsky et al., 2009; Brown et al., 2010). We identified only three isolates in East Africa carrying I164L. A deeper understanding of the evolutionary history of this gene in Africa will require extensive historical data from samples taken around the time of the initial rollout of SP.

The distribution of the *dhps* alleles and haplotypes was more heterogeneous than for *dhfr*. This heterogeneity highlights the differing evolutionary impact of SP use between regions, which should be considered in ongoing SP interventions. A combination of I431V, S436A, A437G, and A613S was dominant in West and Central Africa, while A437G, K540E and A581G were more common in East and North-East Africa. This supports the theory of multiple independent emergences of *dhps* mutant alleles in different African regions (Pearce *et al*., 2009). Within West Africa, the increasing prevalence of A437G in the Gambia seems to have “stalled” in 2016 at around 60%. Since 2016, A437G has been increasing and reached higher prevalence levels in Mali and Ghana. Such difference between countries on the western Atlantic coast and countries in the middle of West Africa has been shown for chloroquine resistance associated alleles (MalariaGEN *et al*., 2023). Assessment of the *crt* K76T locus showed that, between 2013 – 2016, parasites in Ghana, Mali, and Côte d’Ivoire had reverted almost entirely to the wildtype, while the mutant types remained at higher prevalence in Gambia, Senegal and Mauritania. Thus, the local environment and possibly the history and scale up of SP-based interventions may play an important role in the dynamics of SP resistance alleles. Beyond this, there could be other effects impacting SP resistance haplotypes such as the host genetic background in the vector and the human host (Mharakurwa *et al*., 2011). In Mali and Ghana, the prevalence of S436A has been declining while the rarer A613S mutation has recently emerged and spread. Combined with the emergence of I431V and the low but increasing frequencies of K540E and A581G, these additional mutations could seriously impact the efficacy of SP (Oguike *et al*., 2016).

Given the concurrent increase of K540E and A437G in Kenya, it is perhaps surprising that K540E did not spread in West Africa, where A437G is highly prevalent (>60%). In West Africa, a more commonly co-occuring mutation with A437G is S436A possibly because it provides a fitter phenotype. K540E was rare in Madagascar but common in the nearest continental mainland. Investigation of other highly differentiated SNPs between East Africa and Madagascar revealed several other loci outside of *dhfr* and *dhps* that may be involved in drug resistance. For example, the S258L mutation within *aat1* is associated with chloroquine resistance, with a fitness cost to the parasite (Amambua-Ngwa *et al*., 2023). The S258L mutation in *aat1* was also shown to have increased to near-fixation in The Gambia over a similar time span to this study (the years 1984 - 2014). Similarly, the SNPs 3’ of *mdr1* are notable as *mdr1* has speculatively been linked to amodiaquine resistance (Holmgren *et al*., 2007; Humphreys *et al*., 2007). Further studies could make use of the Pf7 dataset to perform a continent-wide assessment of these loci. The exploration of loci beyond *dhfr* and *dhps* was possible thanks to whole-genome sequencing data. These also enabled the calculation of full-gene haplotypes for SP resistance genes, which revealed important variation in the reliability of certain proxy markers between different African regions. Further related work using whole genome data could include the investigation of intermediate haplotypes between the wild type and SP-resistance haplotypes for *dhps* and *dhfr*. For example, many studies have compared *dhfr* NCS vs IRN haplotypes and showed similar MS calls in the latter but not the former (Nair *et al*., 2003; Roper *et al*., 2003; Maïga *et al*., 2007). Using available WGS data to generate and then compare microsatellite loci around *dhfr* NCN, ICN, and NRN haplotypes could provide us with a more detailed picture on the evolution of SP resistance.

We provided an insight to SP resistance haplotype frequencies across 22 countries, but we were limited in our ability to describe temporal trends within these locations due to a lack of consistent sampling across the study period. Longitudinal data from four countries enabled us to highlight differing mutation dynamics within West Africa and between East and West Africa. Increasing the number of countries and years for which sufficient data exists requires longitudinal sampling. These data would increase the specificity of conclusions drawn about SP evolution in Africa, which is essential to ensure the continued efficacy of SP-based interventions such as SMC. In the absence of other openly available time series data from Africa, databases such as those of the Worldwide Antimalarial Resistance Network (WWARN - https://www.iddo.org/wwarn) could be explored. For example, since 2000, WWARN publishes information on the *dhfr* and *dhps* mutations in their SP Molecular Surveyor, collated across 77 countries. However, replicating the analyses of this study would not be possible with WWARN data because full gene haplotypes are not always represented. For example, *dhfr* multi-mutant haplotypes with I164L or the joint *dhps* 436 and 437 haplotypes are not included. Moving towards whole genome sequencing could increase the availability of shared *dhfr, dhps* haplotypes within such databases.

This study has highlighted the persistence and regional variability of SP resistance markers in *P. falciparum* across Africa, possibly driven by the continued use of SP for chemoprevention. The recent, near-fixation of the *dhfr* triple mutant haplotype in several countries and the wide diversity of *dhps* haplotypes across the continent underscore the ongoing selection pressure from SP use. The distinct regional patterns of allele frequencies illustrate the complex dynamics of SP resistance, shaped by local ecological factors, migration, and past antimalarial treatment policies. We have highlighted several avenues for further exploration, including tracing the evolutionary origins of the *dhfr* double mutant haplotypes and the assessment of several SNPs beyond *dhfr* and *dhps* in regions beyond East Africa. To ensure the future effectiveness of SP-based interventions, continued molecular surveillance is essential. Whole-genome sequencing can play an important part in this surveillance by offering a view of resistance evolution beyond established markers. Additionally, moving towards longitudinal sampling would enhance our understanding of changes in SP resistance dynamics over time, informing targeted, region-specific strategies to combat malaria in Africa.

## Methods

Full gene haplotypes, generated from concatenated exon sequences, and their summaries across populations were generated using a custom Python script. For a temporal analysis of mutation frequencies, we investigated variation at four *dhfr* loci linked to SP resistance: 51, 59, 108, and 164 and six *dhps* loci: 431, 436, 437, 540, 581, and 613. We used the allele frequency for all *dhfr* and *dhps* markers. There are multiple ways in which mutation frequencies can be calculated (see Box), but here we excluded from further analysis any sample which had a heterozygous mutation at any position within the coding region of the gene and calculated frequencies from homozygous haplotypes only. Custom Python scripts were used to compare marker allele frequencies between countries for which we had sufficient data and time points with at least 25 homozygous samples: the Gambia, Mali, Ghana, and Kenya.

Maps of dhps A437G and K540E allele frequencies across Africa were generated using the mapMutationPrevalence function from the grcMalaria R package (version 2.0.0) (Verschuuren *et al*., 2024). Investigations of genetic dissimilarity between East Africa and Madagascar were performed using a neighbour-joining tree and a genome-wide comparison of F_ST_ and allele frequencies between these population subsets. The neighbour-joining tree was based on the same genetic distance matrix as calculated for Pf7 (MalariaGEN *et al*., 2023), but a subset to include only Malagasy samples and a random subset from each African population using custom R scripts. Each African population (West, Central, Northeast, and East without Madagascar) was subset to 50 random individuals, while maintaining the contributing proportions of each country found in the full dataset. F_ST_ was computed from QC-passed SNPs and samples from Madagascar (n = 24) and from the remainder of East Africa (n = 1,515) using scikit-allel. The same was done for the genomic position encoding the K540E mutation (Pf3D7_08_v3:549993), and values were compared to genome-wide estimates.

To assess the reliability of single markers as proxies for full gene haplotypes, we calculated the proportion samples with each single marker proxy, which also held the correct, associated full gene haplotype. This proportion was denoted as a marker “success rate”. Proxy marker success rates were generated for countries with >15 samples using custom Python scripts.

### Box - Methods of calculating allele frequencies

Because some malaria samples contain parasite clones with different genotypes (sometimes referred to as complexity of infection or CoI), the genotype calls for a mutation can be heterozygous. When calculating frequencies, decisions need to be made about how to treat heterozygous calls. Assume we have *R* homozygous reference calls, *A* homozygous alternative calls and *H* heterozygous calls. One approach that is sometimes used is to calculate the carrier frequency. This is the proportion of samples that carry the alternate allele and is calculated as 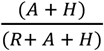. When considering drug-resistance mutations, it is sometimes assumed that if a patient contains both sensitive and resistant parasites, the infection should be considered drug-resistant on the basis that in the presence of the drug, the resistant parasites are likely to out-compete the sensitive parasites, and therefore the carrier frequency is perhaps the most accurate way to estimate the proportions of infections that will be resistant.

Population genetics, however, tends to be more concerned with allele frequencies in the population of all parasites. One unbiased estimate of the allele frequency is to simply ignore heterozygous genotype calls, and calculate the proportion of homozygous alternate calls as a proportion of all homozygous calls, i.e. 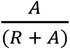. This is the method we have used in this paper. An alternative approach is to weight a heterozygous call as half that of a homozygous alternative call, i.e. calculate allele frequency as 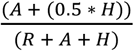.

To take an example from the current study, for samples taken in Kenya in 1995, we see 21 samples with the reference C59 call, 10 samples with the alternative C59R calls and 15 samples with heterozygous calls, so here *R* = 21, *A* = 10 and *H* = 15. The carrier frequency here, calculated using 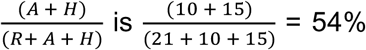 . The allele frequency calculated using 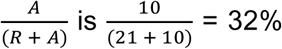. As such there is a difference of 22% between the allele and carrier frequencies.

In Box Figure A we see the differences between allele and carrier frequencies for all the frequencies calculated in this study. In some cases, the values are very close, whereas in other cases, particularly for high minor allele frequencies, the differences can be substantial. Box Figure B shows a histogram of these differences, and it can be seen that many frequencies differ by more than 10%. The difference is particularly pronounced in high-frequency mutations (Box Figure C).

In this study, we have chosen to calculate allele frequencies using homozygous calls only, i.e. using 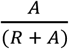.

**Box figure:**
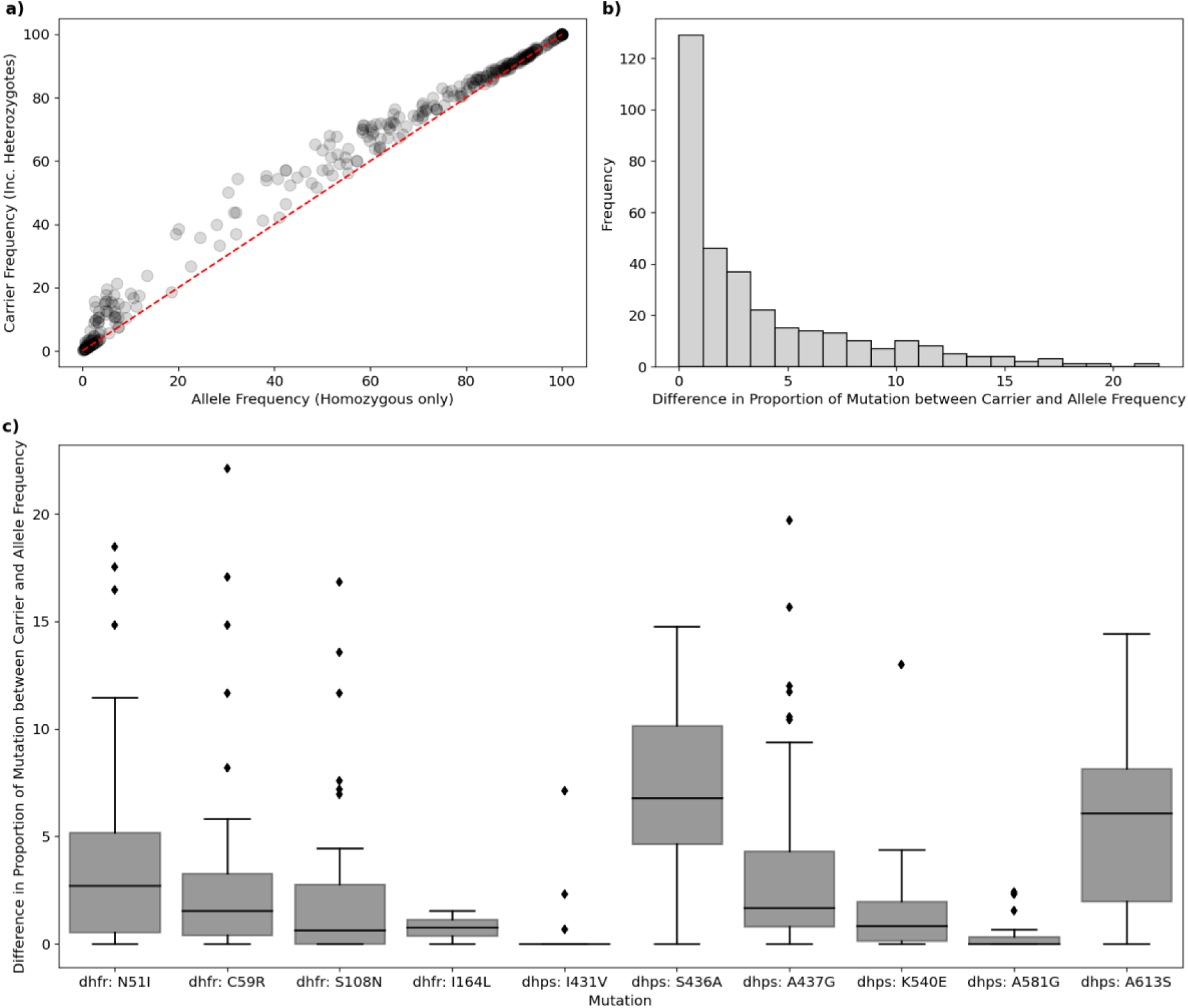
Differences between allele and carrier frequencies. The absolute difference in the carrier frequency and allele frequency calculated for mutations at key drug resistance genes *dhfr* and *dhps*. Data is shown for all Pf7 samples from the Gambia, Mali, Ghana or Kenya, and only for locations and years where the denominator of allele or carrier frequency was at least 25 samples. (a) Identity of carrier vs allele Frequency (b) Histogram of absolute differences between allele and carrier frequencies (c) Differences between frequency calculations by mutation.

