## Supplementary Table S1 for "Disparate co-evolution and prevalence of sulfadoxine and pyrimethamine resistance alleles and haplotypes at *dhfr* and *dhps* genes across Africa"

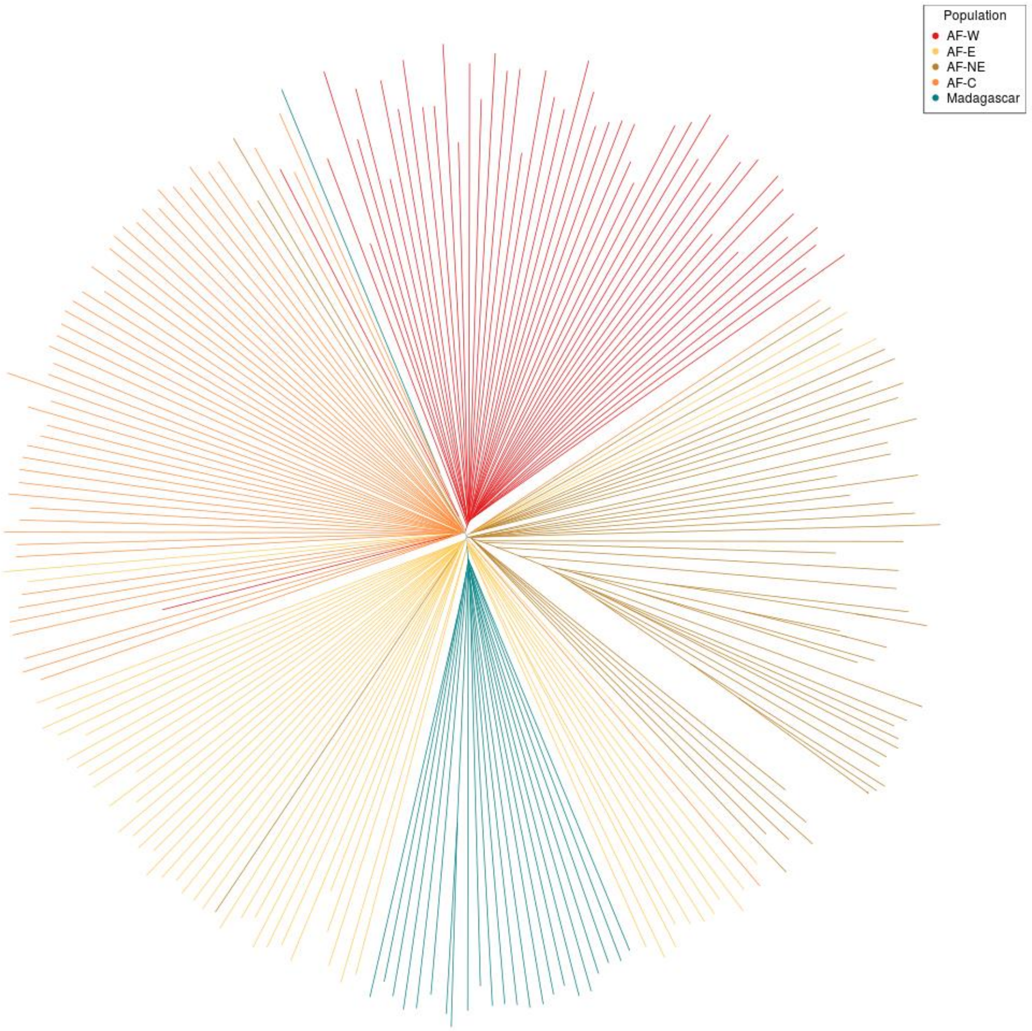


**Supplementary Figure S1**. Genome-wide unrooted neighbour-joining tree between mainland African populations and Madagascar. A random sample of n = 50 QC+ samples was selected for each population: AF-W (west Africa, red), AF-C (central Africa, orange), AF-NE (northeast Africa, brown), and AF-E (east Africa, gold, excluding Madagascar). Madagascar is highlighted in teal and contains n = 24 QC+ samples. Population structure is evident between all populations, but Madagascar is most similar to AF-E.

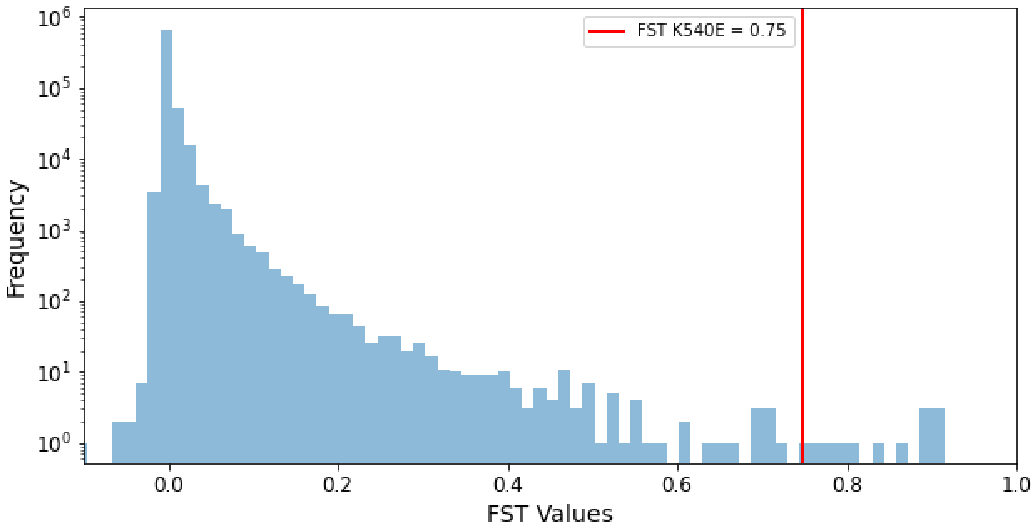


**Supplementary Figure S2**. $F_{ST}$ values for 763,535 genome-wide QC-passed SNPs between Madagascar (n = 24) and the remainder of mainland East Africa (n = 1,515). The $F_{ST}$ value for the nucleotide mutation encoding *dhps* K540E is highlighted in red. Frequency is shown on a log scale.

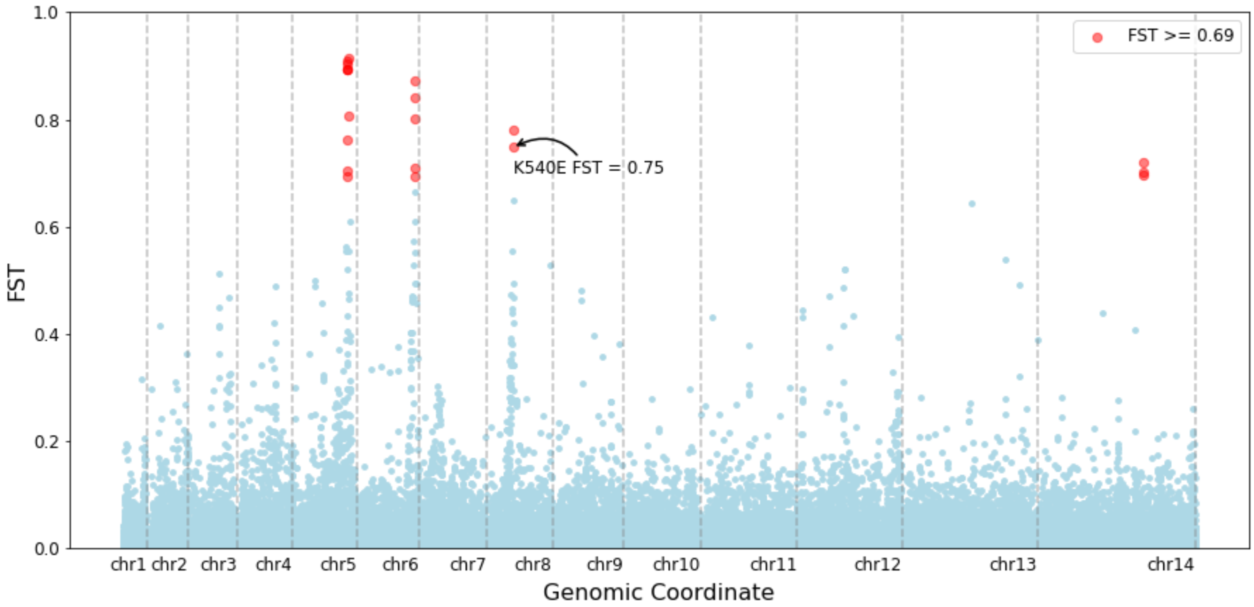


**Supplementary Figure S3**. $F_{ST}$ values for 763,535 genome-wide QC-passed SNPs between Madagascar (n = 24) and the remainder of mainland east Africa (n = 1,515). The 20 most differentiated SNPs ($F_{ST} > 0.69$) are highlighted, including the position encoding *dhps* K540E.

**Supplementary Table S1.** *dhfr* haplotype frequencies by population**.** Haplotype gives the amino acid changes with respect to the 3D7 reference genome, which here is assumed to be the wild type (WT) sequence. Table shows numbers of samples with a homozygous call with each haplotype. Column headings show populations as defined in Pf7: AF-W=Africa-West, AF-C=Africa-Central, AF-NE=Africa-Northeast, AF-E=Africa East. See Pf7 manuscript for further details (MalariaGEN *et al.*, 2023).

| Haplotype | Number of samples | | | | |
|---|---|---|---|---|---|
| | AF-W | AF-C | AF-NE | AF-E | Total |
| **N51I/C59R/S108N** | 3,905 | 373 | 100 | 1,085 | 5,463 |
| **WT** | 661 | 2 | 3 | 47 | 713 |
| **C59R/S108N** | 260 | 3 | 1 | 75 | 339 |
| **N51I/S108N** | 52 | 59 | 45 | 99 | 255 |
| **S108N** | 17 | 2 | 2 | 4 | 25 |
| **N51I** | 0 | 0 | 0 | 2 | 2 |
| **N51I/C59R/S108N/I164L** | 0 | 0 | 0 | 2 | 2 |
| **N157K** | 2 | 0 | 0 | 0 | 2 |
| **C59R** | 1 | 0 | 0 | 0 | 1 |
| **N51I/S108N/I164L** | 0 | 0 | 1 | 0 | 1 |
| **K256R** | 1 | 0 | 0 | 0 | 1 |
| **N51I/C59R/S108N/D414N** | 1 | 0 | 0 | 0 | 1 |
| **N51I/C59R/S108N/D425N** | 1 | 0 | 0 | 0 | 1 |
| **M337K** | 1 | 0 | 0 | 0 | 1 |
| **N51I/C59R/P93S/S108N** | 1 | 0 | 0 | 0 | 1 |
| **T130N** | 1 | 0 | 0 | 0 | 1 |
| **S306F** | 0 | 0 | 0 | 1 | 1 |
| **N90Y** | 1 | 0 | 0 | 0 | 1 |
| **N51I/S108N/S306F** | 0 | 0 | 0 | 1 | 1 |
| **Total** | 4,905 | 439 | 152 | 1,316 | 6,812 |

**Supplementary Table S2.** *dhps* haplotype frequencies by population**.** Haplotype gives the amino acid changes with respect to what is considered to be the wild type (WT) haplotype. Note that in this case, the 3D7 reference genome is not considered to be wild type, as the ancestral allele at amino acid 437 is thought to be alanine (A) where 3D7 has glycine (G). Table shows numbers of samples with a homozygous call with each haplotype. Column headings show populations as defined in Pf7: AF-W=Africa-West, AF-C=Africa-Central, AF-NE=Africa-Northeast, AF-E=Africa East. See Pf7 manuscript for further details (MalariaGEN *et al.*, 2023).

| Haplotype | Number of samples | | | | |
|---|---|---|---|---|---|
| | AF-W | AF-C | AF-NE | AF-E | Total |
| **A437G** | 1,770 | 346 | 2 | 10 | 2,128 |
| **A437G/K540E** | 34 | 23 | 91 | 907 | 1,055 |
| **S436A/A437G** | 943 | 19 | 0 | 1 | 963 |
| **S436A** | 636 | 5 | 3 | 21 | 665 |
| **WT** | 279 | 12 | 20 | 150 | 461 |
| **S436A/A437G/A613S** | 202 | 0 | 0 | 0 | 202 |
| **A437G/K540E/A581G** | 0 | 11 | 12 | 134 | 157 |
| **I431V/S436A/A437G/A581G/A613S** | 72 | 4 | 0 | 0 | 76 |
| **A437G/I484T** | 27 | 0 | 0 | 0 | 27 |
| **E189Q/S436A** | 24 | 0 | 0 | 0 | 24 |
| **I431V/S436A/A437G** | 20 | 0 | 0 | 0 | 20 |
| **S436F/A613S** | 16 | 0 | 0 | 1 | 17 |
| **S152I/A437G** | 16 | 0 | 0 | 0 | 16 |
| **S436C** | 16 | 0 | 0 | 0 | 16 |
| **S436Y/A613S** | 13 | 0 | 0 | 0 | 13 |
| **S436H/A437G/K540E** | 0 | 0 | 7 | 0 | 7 |
| **E189Q/A437G** | 7 | 0 | 0 | 0 | 7 |
| **D280N** | 7 | 0 | 0 | 0 | 7 |
| **A437G/A581G** | 0 | 0 | 0 | 4 | 4 |
| **I431V/S436A/A437G/A613S** | 4 | 0 | 0 | 0 | 4 |
| **A437G/A613S** | 4 | 0 | 0 | 0 | 4 |
| **S436A/A437G/A581G/A613S** | 3 | 0 | 0 | 0 | 3 |
| **R243K** | 3 | 0 | 0 | 0 | 3 |
| **S436A/A613S** | 2 | 0 | 0 | 0 | 2 |
| **V471L** | 2 | 0 | 0 | 0 | 2 |
| **L22F/S436A** | 2 | 0 | 0 | 0 | 2 |
| **R243S/S436A** | 2 | 0 | 0 | 0 | 2 |
| **N13K** | 1 | 0 | 0 | 0 | 1 |
| **R243S** | 1 | 0 | 0 | 0 | 1 |
| **S436A/A437G/K540E** | 0 | 0 | 0 | 1 | 1 |
| **R27K** | 1 | 0 | 0 | 0 | 1 |
| **R78H** | 1 | 0 | 0 | 0 | 1 |
| **S10F** | 0 | 0 | 0 | 1 | 1 |

| | | | | | |
|---|---|---|---|---|---|
| **S436H** | 0 | 0 | 0 | 1 | 1 |
| **L166M** | 0 | 0 | 0 | 1 | 1 |
| **S436A/A672G** | 1 | 0 | 0 | 0 | 1 |
| **L263I** | 1 | 0 | 0 | 0 | 1 |
| **S436A/A581G/A613S** | 1 | 0 | 0 | 0 | 1 |
| **G117E** | 1 | 0 | 0 | 0 | 1 |
| **E189Q/S436A/A437G** | 1 | 0 | 0 | 0 | 1 |
| **K106I/A437G/K540E** | 1 | 0 | 0 | 0 | 1 |
| **D105H** | 1 | 0 | 0 | 0 | 1 |
| **I277T/S436A/A437G** | 1 | 0 | 0 | 0 | 1 |
| **I277L** | 1 | 0 | 0 | 0 | 1 |
| **K540E** | 0 | 0 | 0 | 1 | 1 |
| **D311G/A437G** | 1 | 0 | 0 | 0 | 1 |
| **D250V** | 1 | 0 | 0 | 0 | 1 |
| **E189Q** | 1 | 0 | 0 | 0 | 1 |
| **C351Y/S436A/A437G** | 0 | 1 | 0 | 0 | 1 |
| **E71Q** | 0 | 0 | 0 | 1 | 1 |
| **A437G/I441M** | 1 | 0 | 0 | 0 | 1 |
| **Total** | 4,121 | 421 | 135 | 1,234 | 5,911 |

**Supplementary Table S3.** Single mutations proposed as markers for multi-gene haplotypes.

| **Maker Name** | **Marker** | **Multi Mutant Haplotype** |
|---|---|---|
| Triple | *dhfr* C59R | *dhfr* N51I / C59R / S108N |
| Quintuple | *dhfr* C59R and *dhps* K540E | *dhfr* N51I / C59R / S108N, *dhps* A437G / K540E |
| Sextuple | *dhps* A581G | *dhfr* N51I / C59R / S108N, *dhps* A437G / K540E / A581G |
